# Modeling immunoglobulin light chain amyloidosis in *Caenorhabditis elegans*

**DOI:** 10.1101/2024.07.05.602215

**Authors:** Margherita Romeo, Maria Monica Barzago, Alessandro Corbelli, Silvia Maglioni, Natascia Ventura, Carmina Natale, Andrea Conz, Mario Salmona, Giovanni Palladini, Mario Nuvolone, Fabio Fiordaliso, Giampaolo Merlini, Luisa Diomede

## Abstract

Cardiomyopathy determines the prognosis of patients with immunoglobulin light chain (AL) amyloidosis, a rare systemic disease caused by the misfolding and deposition of monoclonal light chains (LCs). The reasons underlining their cardiac tropism remain unknown, and an animal model recapitulating the main pathological features of AL amyloidosis is instrumental. Taking advantage of the similarities between the vertebrate heart and *C. elegans*’ pharynx, we developed a new transgenic nematode expressing a human amyloidogenic λ LC whose sequence was deduced from a patient suffering from AL amyloidosis with cardiac involvement (MNH). Strains expressing a non-amyloidogenic LC (MNM) or the empty vector only (MNV) were generated as controls. At variance with controls, LCs expressed in the body-wall muscle of MNH worms formed native soluble dimeric assemblies, which were secreted and reached different organs, including the pharynx. Noteworthy, MNH worms exerted a pharyngeal impairment resembling cardiac functional impairment occurring in patients with AL, accompanied by increased radical oxygen species production and tissue ultrastructural damage. This new animal model can allow the elucidation of the mechanisms underlying the cardiac-specific tropism occurring in AL amyloidosis, providing innovative insights into the pathophysiology.

## Introduction

Immunoglobulin light chain (AL) amyloidosis is a severe protein misfolding disease caused by a B cell or a plasma cell clone, resulting in the overproduction of unstable monoclonal light chains (LCs) [1, 2]. After secretion into the bloodstream, LCs are transported to target organs, accumulating and exerting their toxic effects. Among the organs involved, the heart is one of the most frequently damaged and the one that causes the onset of rapidly progressive and fatal cardiomyopathies, determining patients’ survival [2, 3]. Notably, cardiotoxicity is the leading cause of death in AL patients independently from the organs involved since approximately 80-90% die from heart failure or fatal arrhythmias [4–7].

Today, the mechanisms underlying the tissue-specific cardiac targeting and damage of LCs are still undetermined, making it challenging to develop new pharmacological approaches to reduce the mortality of patients suffering from AL amyloidosis [7–11]. Numerous attempts have been made to decipher the onset and progression of the AL pathological process, from amyloidogenic LC secretion to fibrillar deposit formation and organ tropism. Information on the general steps leading LCs to misfold and aggregate has been obtained from i*n vitro* studies [12–19]. Experiments conducted with cardiomyocytes and cardiac fibroblasts provided advances in the mechanism of LC toxicity, showing that soluble oligomers and amyloidogenic deposits are both key players [15, 20–23]. While soluble oligomers reduce cell viability through aberrant interactions with critical cellular components and generation of radical oxygen species (ROS), resulting in mitochondria damage [22], the amyloid burden can contribute to organ damage by mechanical constraint and alteration of the tissue architecture [2, 23].

The effort made over the years to develop adequate animal models of AL has also been particularly intense [24]. In zebrafish, the injection of cardiotoxic LC resulted in cardiac dysfunction, cell death, and early mortality [3, 25]. Rodents proved to be refractory to LC cardiotoxicity [24–28]. The nematode *Caenorhabditis elegans (C. elegans)* has become a major experimental organism with applications to many biomedical research areas. This model organism has spearheaded aging research and is widely used in experimental cardiology [29] since its rhythmically pumping pharynx is considered an ortholog of the vertebrate heart. The worm pharynx is similar to the heart. It is a tubular muscular pump primarily composed of smooth muscle tissue and accessory neurons, capable of self-stimulation and autonomous generation of the depolarising signal independent of extra-organ neural stimulation [30–34]. Contraction of the *C. elegans* pharynx is continuous, involuntary, and persistent throughout the worm’s entire life in a manner that is strictly comparable with the one of a more evolved heart [33, 34].

This model was exploited to investigate the mechanism of cardiac toxicity by administering Bence Jones or recombinant LCs, demonstrating that only proteins from patients with amyloid cardiomyopathy damaged the pharynx of worms [21]. In the present study, we developed a new *C. elegans*-based AL animal model by generating a transgenic MNH strain constitutively expressing a human amyloidogenic λ LC whose sequence was deduced from a patient suffering from AL with cardiac involvement in the body-wall muscle cells. To this end, we used the *mos1*-mediated Single Copy Insertion (MosSCI) method to insert genetic cargo at defined locations in the worm’s genome [35]. Two additional strains were developed as controls: the MNM strain expressing a non-amyloidogenic LC whose sequence was deduced from a patient affected by multiple myeloma with no evidence of AL amyloidosis, and the MNV expressing the empty vector only. The findings indicate that the expression of the cardiotoxic amyloidogenic LC in the body-wall muscle of MNH worms generated a native soluble dimeric protein, which can be secreted, as observed in AL patients. These LCs can reach various *C. elegans* organs but cause a specific pharyngeal impairment, increased superoxide production, and ultrastructural damage that did not occur in MNM and MNV worms.

These new *C. elegans* models offer a practical, informative approach to investigating the mechanisms underlying the tissue-specific tropism of LCs. They provide new insights into the pathophysiology of AL amyloidosis and represent a valuable tool for preclinical studies.

## Materials and methods

### Transgenic C. elegans strains

All *C. elegans* strains were cultured and handled using standard breeding conditions. Experiments were performed at 20°C on standard Nematode Growth Media (NGM) seeded with OP50 *Escherichia coli* (The Caenorhabditis Genetics Center, GCG, Minnesota, USA). To model AL amyloidosis, a new transgenic *C. elegans* strain was generated by Invivo Biosystems (Eugene, USA) through the MosSCI method, allowing the insertion of a single copy of a transgene into chromosome II of the worm [35]. The *C. elegans* strain was engineered to express the human amyloidogenic λ LC H7 constitutively in the body-wall muscle cells under *myo-3* promoter (MNH) [21] (**Supplementary Table 1**). The H7 sequence inserted was deduced from the cDNA isolated from the bone marrow cells of an AL patient with cardiac involvement [51]. As controls, a strain constitutively expressing under *myo-3* promoter the human non-amyloidogenic LC M7 whose sequence was deduced from the cDNA of a patient with multiple myeloma [52] (MNM), and a strain expressing the empty vector alone (MNV) were developed (**Supplementary Table 1**). The two LCs, H7 and M7, were expressed in worms with specific secretion sequences. The full-length H7 and M7 protein sequences are reported in **Supplementary Fig. 1**. Transgenics selection was done by screening strains by Invivo Biosystems by PCR for single-copy insertion at the *mos1* locus. A further transgenic strain was generated by Invivo Biosystems inserting the mCherry DNA sequence at the C-terminus of the H7 sequence expressed by MNH using CRISPR-site direct mutagenesis (MNH::mCherry) (**Supplementary Table 1**). The relative levels of H7 and M7 genes expressed by MNH and MNM worms were determined by quantitative PCR (Q-PCR) (**Supplementary Information and Supplementary Table 2).**

### Pharyngeal and Motility Assays

The nematodes were synchronized by egg-laying and cultured on NGM plates seeded with *E. coli* OP50 for food at 20°C. From the first to the eighth day of adulthood, the pharyngeal pumping rate, resulting from counting the number of times the terminal bulb of the pharynx contracts in a 1-min interval (pumps/min) [21], and the motility test, based on counting the number of body movements in liquid (body bending/min) in a 1-min interval [53], were performed.

### Electropharyngeogram

The nematodes were synchronized by egg-laying and cultured on NGM plates seeded with *E. coli* OP50 for food at 20°C. On the first day of adulthood, they were collected with M9, settled by gravity, and washed with M9 to eliminate bacteria. They were transferred in a reaction tube, washed three times with M9, and incubated in a 10 mM serotonin solution (Sigma-Aldrich) for 30 min. The ScreenChipTM System (InVivo Biosystems) was employed to record the voltage changes caused by the contraction of the pharynx in real-time, producing the so-called electropharyngeogram (EPG)[54]. Worms were loaded on the ScreenChip SC40 (InVivo Biosystems) with a syringe (0.01 ml – 1 ml), and the EPG of single worms was recorded for about 2 minutes. Only worms that showed pumping activity were recorded, while those without pumping activity were discarded. The software programs NemAcquire 2.1 and NemAnalysis 0.2 (https://invivobiosystems.com/product-category/instruments/screenchip-system-software/) were used for recording and analysis, respectively. The following parameters were measured: pump frequency, spike amplitude ratios, pump duration, and interpump interval (IPI) [54].

### Transmission electron microscopy

Age-synchronized worms, on the first day of adulthood, were picked, and a 2% glutaraldehyde solution containing 4% paraformaldehyde in 0.12 M phosphate buffer, pH 7.4, was introduced in the head by microinjection. Worms were cut open at the level of the second bulb of the pharynx to improve access to the fixative and then left in the fixative for 1 week at room temperature. After post-fixation in a solution of 1% OsO4 and 1.5% ferrocyanide in 0.12 M cacodylate buffer (ferrocyanide-reduced OsO4) at 4°C for 2 h, samples were incubated in 0.5% thiocarbohydrazide in cacodylate buffer 0.12 M for 30 min at room temperature and finally in 1% OsO4 in 0.12 M cacodylate buffer for 1 h at 4°C. The pharynx was then dehydrated in a graded series of ethanol and then infiltrated at first in a graded series of epon (Epon 812, Merk Life Science S.r.l., Milano, Italy) in acetone, and finally embedded in 100% of epon, followed by polymerization at 60°C (Leica Microsystems GmbH, Wetzlar, Germany) for 72 h.

From each sample, one semithin (1 μm) section was cut with a Leica EM UC6 ultramicrotome and mounted on glass slides for light microscopic inspection. Ultrathin (60–80 nm thick) sections of areas of interest were obtained, counterstained with uranyl acetate and lead citrate, and examined with an Energy Filter TEM (ZEISS LIBRA® 120) equipped with a YAG scintillator slow scan CCD camera.

### Pharmacological studies

Pharmacological studies were performed as previously described by our group [21, 22]. On the first day of adulthood, age-synchronized MNH and MNV worms were collected, centrifuged (100 x *g* for 3 minutes), and washed twice with M9 buffer. The nematodes (100 worms/100 μl) were treated for 2 h in 1.5 ml tubes with 0-1 mM ascorbic acid (Sigma-Aldrich), 0-200 μM doxycycline hydrochloride (Sigma-Aldrich), and 0-2 nM PBT2 (Prana Biotechnology Ltd, Parkville, Australia) [22] Using the same protocol, worms were treated for 2 h with 10 mM PBS, pH 7.4, (100 worms/100μl) as control (Vehicle). At the end of incubation, worms were transferred to NGM plates seeded with OP50 bacteria and spotted with 0-1 mM ascorbic acid, 0-200 μM doxycycline, or 0-2 nM PBT2. The pumping rate was measured 20 h later, as described above.

### Statistical analysis

No randomization was required for *C. elegans* experiments. All evaluations were done blind to the sample identity and treatment group. The data were analyzed using GraphPad Prism 9.4.1 software (CA, USA) by Student’s t-test, one-way or two-way ANOVA, and Bonferroni’s *post hoc* test. IC50 values were determined using the same software. A p-value <0.05 was considered significant. For lifespan and healthspan studies, the number of dead and censored animals was used for survival analysis in OASIS 2 [55]. The p-values were calculated using the log rank and Bonferroni’s *post hoc* test between the pooled populations of animals.

## Results

### Generation of transgenic *C. elegans* strains

To develop a new animal model of AL amyloidosis, we generated a transgenic MNH *C. elegans* strain constitutively expressing the human amyloidogenic H7 LC with its specific secretion sequence deduced from an AL patient with cardiac involvement under the body-wall muscle-specific promoter *myo-3* (**Fig. 1a; Supplementary Fig. 1**).

**Fig. 1.**
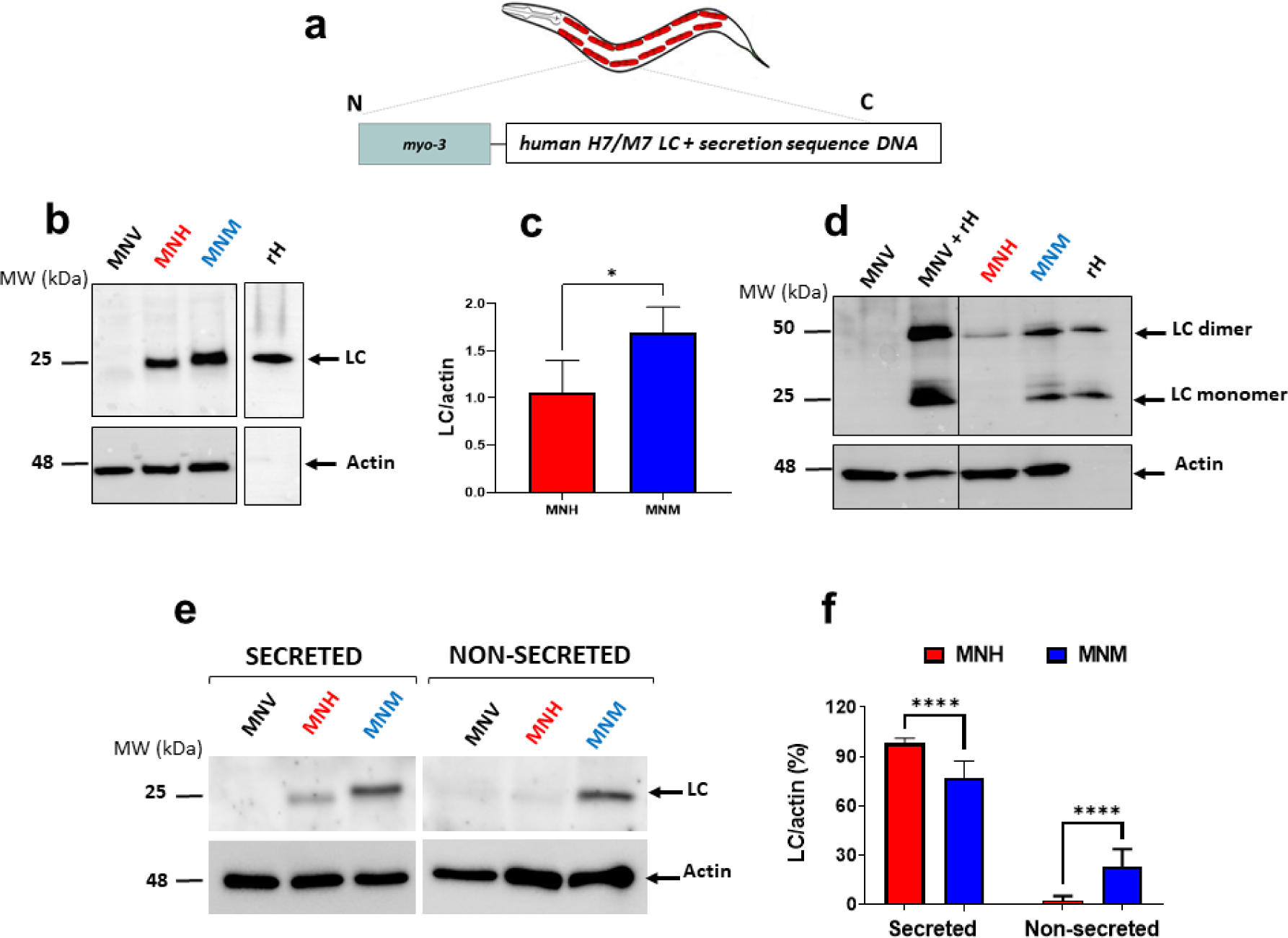
Characterization of the transgenic *C. elegans* strains. (**a**) Schematic representation of transgenic *C. elegans* transgenes generated in this study. The amyloidogenic cardiotoxic (H7) and non-amyloidogenic (M7) LCs were expressed under the body-wall muscle-specific promoter *myo-3* to generate MNH and MNM transgenic *C. elegans* strains. The human-specific secretion was included in each strain to enable LC to be secreted. (**b, d)** Western blots, representative of 4 biological replicates performed in (**b**) reducing and (**d**) non-reducing conditions of total LCs expressed by worms on the first day of adulthood. Equal amount of proteins (25 µg) were loaded in each gel lane and immunoblotted with anti-human λ total LC or anti-actin antibody. Recombinant H7 LC (rH, 50 ng) was loaded alone or with 25 µg of protein lysate of MNV worms as additional positive controls. (**c**) Quantification of total LCs expressed as the mean volume of the anti-human λ total LC band immunoreactivity of the Western blot in the **(b)** normalized on actin band. Data are mean ± SD (n = 4). (**e**) Representative Western blots of secreted and non-secreted fractions of worm’s lysates on the first day of adulthood. Equal proteins (25 µg) were loaded in each gel lane and immunoblotted with anti-human λ total LC or anti-actin antibody. (**f**) Percentage of secreted and non-secreted LCs (mean ± SD, n = 3). p<0.0001, Student’s t-test.

A strain expressing a human non-amyloidogenic M7 LC and its secretion sequence deduced from a patient with multiple myeloma (MNM) (**Fig. 1a; Supplementary Fig. 1**) and a strain expressing the empty vector only (MNV) under the *myo-3* promoter were generated as controls. Transgenic worms were produced employing the MosSCI technique, which allows the insertion of a single copy of a transgene into chromosome II of the worm. We assessed the LC mRNA and protein levels expressed by the synchronized transgenic strains on the first day of adulthood. MNH worms expressed a significantly lower mRNA LC level than MNM; as expected, no signal was observed for MNV transgenic animals (**Supplementary Fig. 2**). Accordingly, MNH worms produced a significantly lower amount of LC protein than MNM, as indicated by SDS-PAGE analysis under reducing conditions (**Fig. 1b-c**). No signal was detected in MNV worms (**Fig. 1b**), indicating that the antibody employed did not recognize any *C. elegans* endogenous proteins. Western blot analysis performed under non-reducing conditions showed that in MNH worms, LCs are expressed only as dimers. In contrast, in MNM, the proteins are produced both as monomers and dimers (**Fig. 1d**).

Amyloidogenic LCs are efficiently secreted by plasma cells and released into the bloodstream in humans. To evaluate if the LCs produced by MNH worms in the body-wall muscle cells can be secreted too, we performed SDS-PAGE analysis on the secreted and non-secreted fractions obtained from homogenates of transgenic worms on the first day of adulthood (**Fig. 1e**). Higher immunoreactive signals were observed in Western blot obtained from secreted and non-secreted fractions of MNM compared to MNH worms, confirming that the latter produced lower amounts of LCs (**Fig. 1e**). However, the quantification of the immunoreactive bands in the secreted and non-secreted fractions indicated that the percentage of protein secreted by MNM worms was significantly lower than that of the one secreted by MNH worms (76.9 % ± 10.5 and 98.0 % ± 3.1 LC secreted by MNM and MNH, respectively) (**Fig. 1f**). This data indicates that almost all the amyloidogenic LC produced by MNH worms was released in the extracellular space. To confirm this finding, worms’ lysates were sequentially fractionated into high-salt reassembly buffer (RAB) and detergent-soluble radioimmunoprecipitation buffer (RIPA) to separate the soluble extracellular LC from those complexed with the membrane [36] (**Supplementary Fig. 3a**). A higher percentage of RAB-soluble extracellular LC and a lower percentage of protein in the RIPA fraction were found in MNH compared to MNM worms (**Supplementary Fig. 3b**), proving the amyloidogenic LC’s greater availability in the extracellular space.

To follow the fate of the LCs secreted by transgenic worms, we planned to generate strains in which LCs were expressed and linked to the mCherry tag. Only worms expressing the amyloidogenic LC fused with mCherry were obtained (MNH::mCherry) because the expression of non-amyloidogenic protein fused with mCherry was not compatible with the survival of nematodes. Western blotting analysis indicated that transgenic MNH::mCherry worms, on the first day of adulthood, expressed an amount of LCs comparable to that produced by MNH nematodes of the same age (**Supplementary Fig. 4a,b**) and exhibited a similar pharyngeal dysfunction (**Supplementary Fig. 4c**). Confocal analysis performed on MNH::mCherry worms indicated a mCherry signal in the body-wall muscles and coelomocytes (**Fig. 2a**), specific cells with phagocytic activity [37]. This demonstrates that after LCs are synthesized in body-wall muscle cells, they are secreted and then taken up by coelomocytes, accumulating in large vesicles [38] (**Fig. 2b)**. Noteworthy, amyloidogenic LCs can also reach other organs, particularly the pharynx of worms (**Fig. 2a**).

**Fig. 2.**
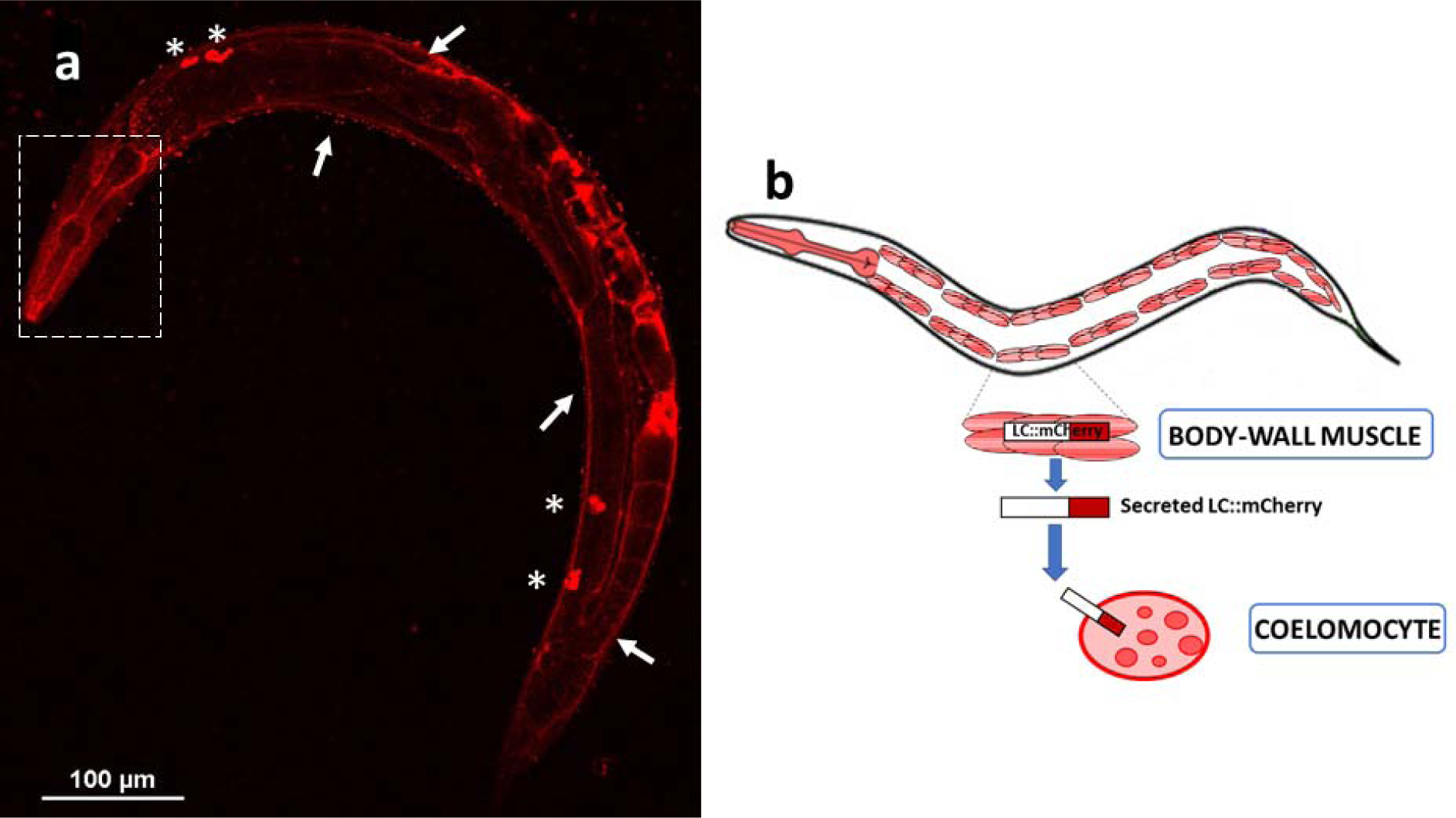
Localization of the amyloidogenic LC in MNH::mCherry worms. (**a**) Representative confocal image of MNH::mCherry worms on the first day of adulthood showing a positive mCherry signal in the body-wall muscle (white arrows) and in coelomocytes (asterisks), proving the uptake of the secreted LC. A positive signal was also detected in the anterior region of the worm, including the pharynx (white inbox). (**b**) A picture representing LC-tagged synthesized in body-wall muscle cells by MNH::mCherry nematodes, their secretion pathway, and the uptake by coelomocytes.

Taken together, these data show that expression of a cardiotoxic amyloidogenic LC in the body-wall muscle of worms resulted in the generation of a soluble dimeric protein which, as observed in AL patients, can be secreted reaching the *C. elegans* “ancestral heart”: the pharynx.

### The expression of the amyloidogenic LC in MNH worms caused a specific pharyngeal dysfunction and structural damage

Experiments were performed to investigate whether the expression of the amyloidogenic LC in MNH worms translated into the onset of specific phenotypic dysfunctions. To this end, various behavioral tests were performed on MNH, MNM, and MNV nematodes. We measured the neuromuscular activity of the transgenic worms by counting the number of movements in a liquid and their pharyngeal motility by scoring the number of pharyngeal contractions in a minute (**Fig. 3a, b**).

**Fig. 3.**
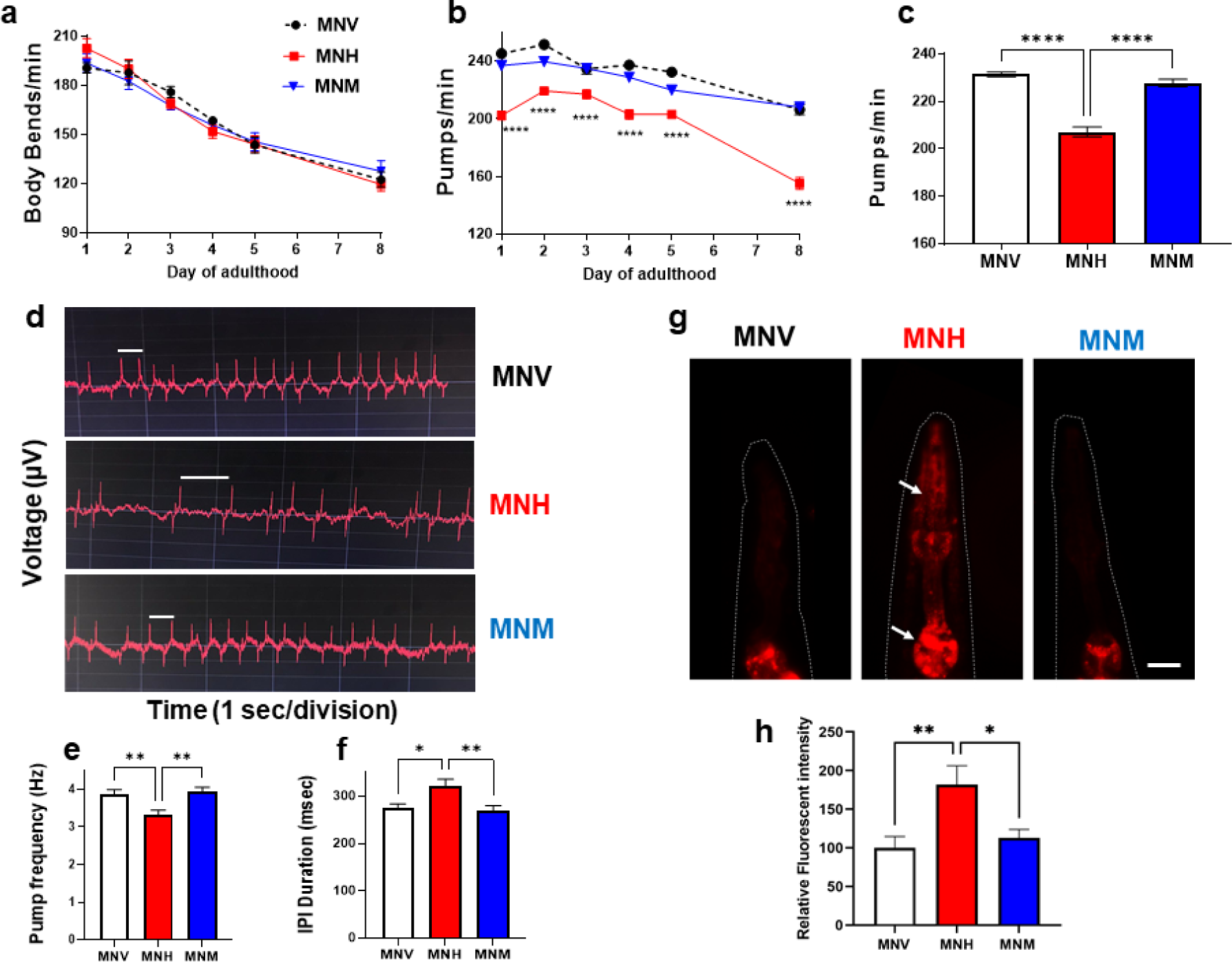
The expression of human cardiotoxic LC in MNH transgenic worms induced a specific pharyngeal impairment. (**a**) Transgenic worms’ motility and (**b**) pharyngeal activity from the first to the eighth day of adulthood. Data are expressed as the mean of (**a**) body bends/min or (**b**) pumps/min ± SEM (n = 20 worms/assay, 3 assays). **** p<0.0001 vs. the corresponding MNV and MNH time point, one-way ANOVA, and Bonferroni’s post hoc test. (**c**) The pharyngeal activity of worms on the first day of adulthood is expressed as pumps/min. Data are the mean ± SEM (n = 40 worms/assay, 4 assays). **** p<0.0001, one-way ANOVA, and Bonferroni’s post hoc test. (**d**) Representative electropharyngeograms worms on the first day of adulthood. The voltage change was plotted against the time. A positive voltage change can be observed as an excitatory spike upon pharynx contraction. The relaxation of the pharynx muscle resulted in an adverse voltage change, the relaxation spike. (**e**) The pump frequency was expressed in Hz. (**f**) The interpump interval (IPI) (i.e., the time between two excitatory spikes, white line) was defined in msec. (**e,f**) Data are the mean ± SEM (n = 30 worms/assay, 3 assays). * p<0.001 and ** p<0.005, one-way ANOVA, and Tukey’s post hoc test. (**g**) Representative images of mitochondrial superoxide production in the pharynx of worms, detected with MitoSOX™ Red. Indicated by red fluorescence, an increase in intracellular superoxide production was observed in MNH worms’ anterior and terminal pharyngeal bulbs (arrows). Scale bar = 50 µm. (**h**) Relative quantification of MitoSOX™ red fluorescence intensity measured by ImageJ software. Data are the mean ± SEM (n = 30 worms/assay, 3 assays). * p<0.001 and ** p<0.005, one-way ANOVA, and Bonferroni’s post hoc test.

In MNH worms, from the first to the eighth day of adulthood, we observed a physiological decline of the neuromuscular activity similar to that of the MNM and MNV control strains (**Fig. 3a**), indicating that, at these ages, the expression of the amyloidogenic LC in the body-wall muscle cells did not cause any specific neuromuscular defect. Similar data was obtained from the evaluation of worms’ healthy aging by scoring the ability of animals to crawl spontaneously or after a manual stimulus (healthspan) (**Supplementary Fig. 5a)**. However, starting from day 12 of age, the amyloidogenic LC’s expression worsted MNH’s ability to move spontaneously compared to MNV (**Supplementary Fig. 5a**). This resulted in a significant reduction of the median health span of MNH nematodes compared to MNV (15.1 ± 0.3 days and 16.1 ± 0.4 days for MNH and MNV, respectively, p= 0.008 log-rank and Bonferroni’s *post hoc* test) (**Supplementary Table 3)** and indicated that the expression of amyloidogenic LC impaired the muscular function of MNH worms during aging. This finding aligns with the knowledge that *C. elegans’* body-wall muscle cells are considered orthologues of the cardiomyocytes [29]. However, the specificity of this effect is difficult to establish because a significant reduction in health span was also observed in MNM worms compared to MNV and MNH (**Supplementary Fig. 5a; Supplementary Table 3**). Moreover, the amyloidogenic and non-amyloidogenic LC expression in MNH and MNM worms significantly reduced their survival compared to MNV (**Supplementary Table 3; Supplementary Fig. 5b**).

In MNH nematodes, we observed a significant impairment of the pharyngeal function, which worsened with age (**Fig. 3b**). This defect was explicitly related to the expression of amyloidogenic LC since no pharyngeal impairment occurred in MNM and MNV worms, in which only a physiological age-dependent decline was observed (**Fig. 3b**). On the first day of adulthood, the pharyngeal activity of MNH worms was ∼ 15% lower than that of MNV and MNM (**Fig. 3c**) and became ∼24 % lower at day 8 (**Fig. 3b**). In addition, electropharyngeograms registered in worms on day 1 of adulthood indicated that MNH worms exhibited a significantly reduced pharyngeal pump frequency and an increase in the interpump interval (IPI) (i.e., the interval between an excitatory spike and the following relaxation spike) compared to MNM and MNV (**Fig. 3d-f**). These data confirmed that the amyloidogenic LC responsible for the onset of amyloidosis with cardiac involvement in AL patients caused a pharyngeal-specific dysfunction when expressed in *C. elegans*. One of the mechanisms underlying the proteotoxicity of amyloidogenic LCs is their ability to increase ROS production, particularly in mitochondria, which we have documented in the *C. elegans* model [21, 22]. To evaluate whether the pharyngeal dysfunction in MNH worms was linked to superoxide production, nematodes were fed with MitoSOX™ Red, able to permeate live cells selectively targeting mitochondria. As shown in **Fig. 3g-h**, in the pharynx of MNH, there was a strong increase in the fluorescent red signal compared to MNV and MNM, indicative of a significant increase in the superoxide production in the mitochondria of animals expressing the amyloidogenic LC. The fluorescent signal observed in the MNH pharynx was similar to that caused in MNV by administering exogenous H_2_O_2_, which is used as a chemical stressor (**Supplementary Fig. 6**).

Ultrastructural studies were then performed to evaluate whether the pharyngeal dysfunction can also be linked to alterations in the organ subcellular compartments, particularly mitochondria, which play a vital role in providing energy for contractile activity. In particular, transmission electron microscopy (TEM) studies were performed on the transverse sections of transgenic animals on the first day of adulthood to analyze the pharynx (**Fig. 4a-d**) and body-wall muscles (**Fig. 4e-h)**. Profound ultrastructural alterations of contractile apparatus were observed in the pharynx of MNH worms, with disruption of the contractile filaments in pharyngeal muscles and mitochondrial damage in pharyngeal muscles and marginal cells (**Fig. 4c**), but not in the pharynx of MNM (**Fig. 4d**) whose morphology was comparable to MNV pharynx (**Fig. 4b**). No ultrastructural damage of myofilaments or mitochondria was observed in the body-wall muscles of MNH (**Fig. 4g**) and MNM at this age (**Fig. 4h**) compared to MNV worms (**Fig. 4f**), even though LCs are expressed in this tissue compartment.

**Fig. 4.**
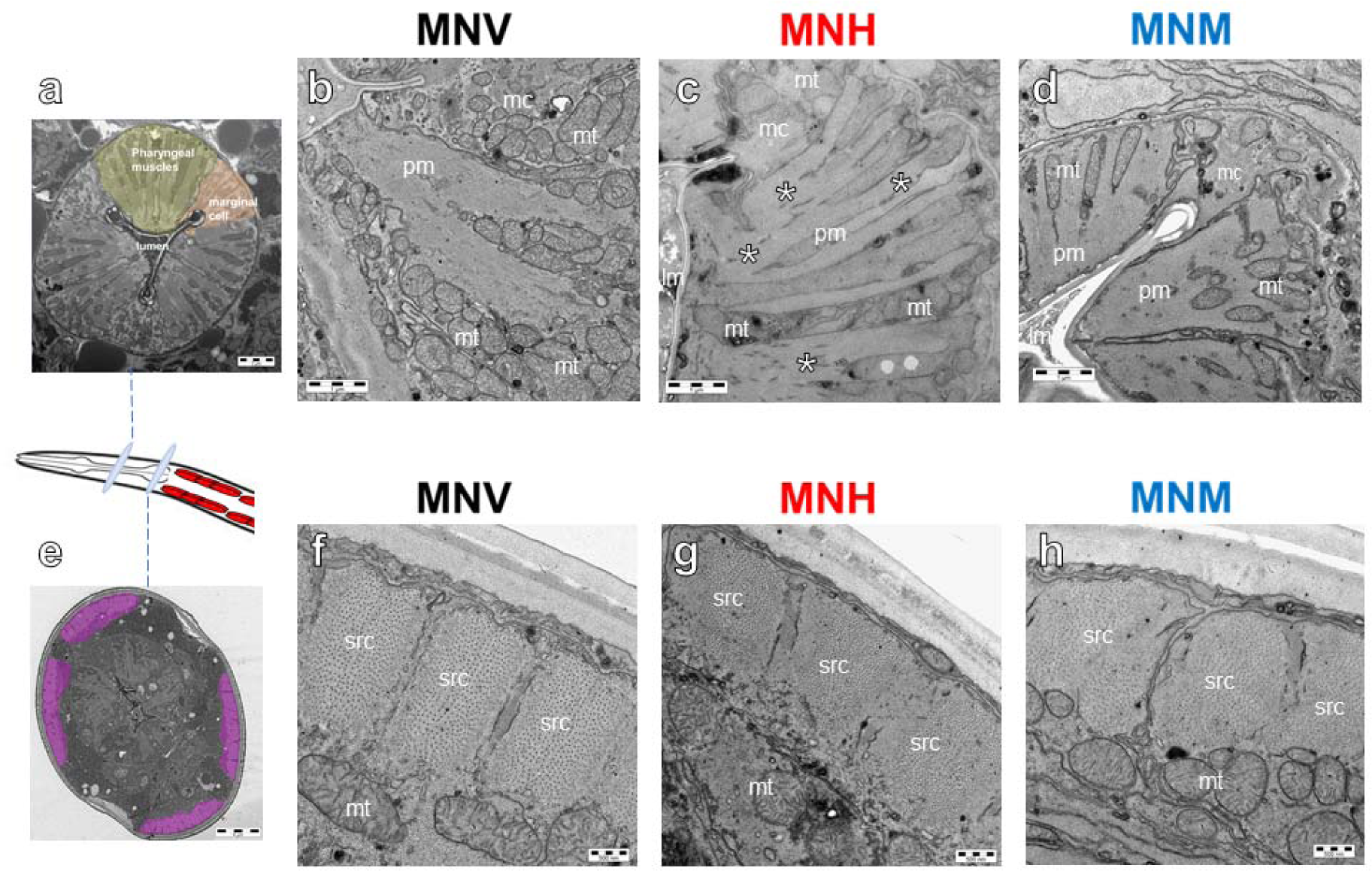
The expression of cardiotoxic LC in MNH worms severely affects pharyngeal ultrastructure. Representative images of the (**a-d**) worm’s pharynx and (**e-h**) body wall muscles obtained from the ultrastructural analysis by TEM in the different strains. (**a**) The transverse section at low magnification above the second bulb of the pharynx showed the characteristic pharyngeal structure with pharyngeal muscles (pm, in green) separated by a marginal cell (mc, in orange), placed at the corner of the pharyngeal lumen (lm), also observed at higher magnification in the pharynx of (**b**) MNV and (**d**) MNM worms. Conversely, the (**c**) MNH pharyngeal ultrastructure was seriously compromised with the presence of amorphous material (asterisks) in place of the radially oriented contractile filaments in pharyngeal muscles. Mitochondrial (mt) internal components in pharyngeal muscles and marginal cells were damaged. (**e**) The transverse section at low magnification of body wall muscles (in purple). No ultrastructural damage of longitudinally oriented myofilaments of sarcomeres (src) or mitochondria was observed in the body wall muscles of (**h**) MNM and (**g**) MNH compared to (**f**) MNV worms.

To investigate whether the pharyngeal dysfunction can be related to the formation of amyloidogenic LC deposits, X-34, a highly fluorescent derivative of Congo red [39], was administered to MNH and MNH::mCherry worms from the first to the fifth day of adulthood [40]. No X-34-positive deposits were observed in the pharynx nor the body-wall muscle at any of the ages considered (data not shown), indicating that the pharyngeal-specific toxicity in MNH worms can be ascribed to the presence in the tissue of soluble LC conformers.

### MNH strain can be used for preclinical studies

We also performed some experiments to explore if the MNH strain can be used as an AL animal model to discover and test novel pharmacological treatments. To this end, we selected ascorbic acid as the prototypic antioxidant already demonstrated to revert the toxicity of cardiotoxic LCs administered to worms thanks to their ability to counteract ROS-induced pharyngeal damage [22]. We also used doxycycline, which reduced the LC aggregation in a transgenic mouse model of AL amyloidosis and the toxicity of LCs administered to *C. elegans* [21, 41]. Based on our previous findings demonstrating the pivotal role of metal ions, particularly copper, in LC-induced toxicity [14, 22], PBT2, an 8-hydroxyquinoline derivative acting as copper/zinc ionophore, was used as a metal-chelating compound [42]. When administered alone, PBT2 permanently blocks ROS production and prevents the toxic effects caused in *C. elegans* by feeding amyloid LCs [22]. We observed that a single administration of all the drugs tested reduced, in a dose-dependent manner, the pharyngeal dysfunction of MNH worms on the first day of adulthood (**Fig. 5a-c**). IC_50_ values in the same order of magnitude were obtained for ascorbic acid (14.38 µM ± 1.42) and doxycycline (9.70 µM ± 1.21) (**Fig. 5a-b**). Starting from 50 µM, doxycycline became toxic, as indicated by its ability to significantly reduce the pharyngeal function of MNV worms (**Fig. 5b**). PBT2 was the most effective compound with an IC_50_ value of 0.09 nM ± 1.1.26 (from 108,000 to 160,000-fold more effective than the other drugs tested) (**Fig. 5c**). When administered to MNH worms at their optimal concentration, ascorbic acid (57 µM), doxycycline (25 µM), and PBT2 (0.5 nM) allowed the full recovery of the pharyngeal defect caused by the amyloidogenic LC expression (**Fig. 5d**). These findings indicate that this strain can provide a useful tool for investigating the efficacy of drugs in protecting against LC-induced tissue damage.

**Fig. 5.**
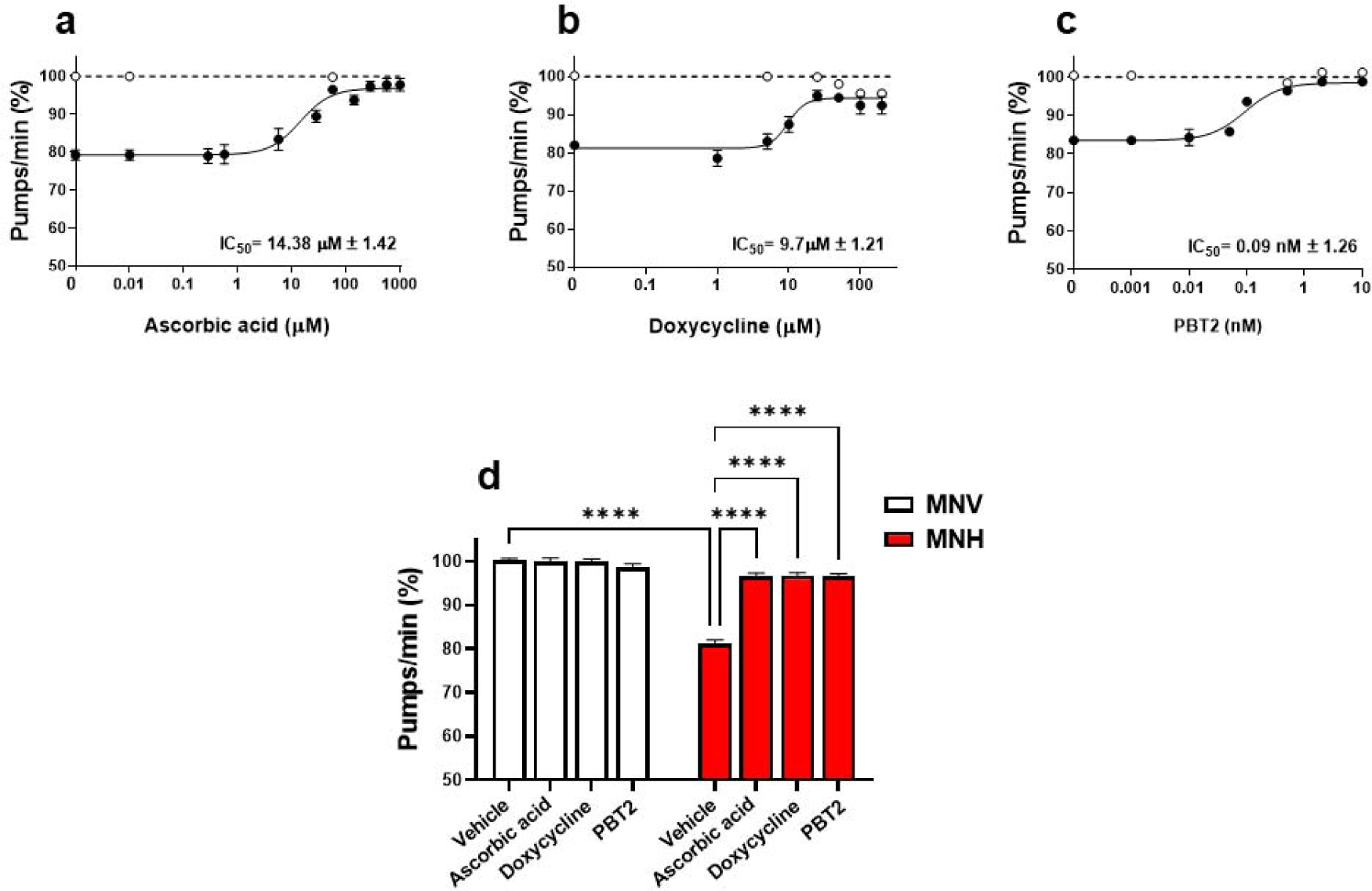
Use of the MNH worms for preclinical studies. Dose-response effect of (**a**) ascorbic acid, (**b**) doxycycline, and (**c**) PBT2 on pharyngeal dysfunction in MNH worms. MNV (white circles) and MNH (black circles) worms on the first day of adulthood were treated for 2 h at room temperature with increasing drug concentrations (100 worms/100 µl). Control MNV worms were treated with 10 mM PBS, pH 7.4 (dotted line). Worms were then plated on NGM seeded with *E. coli,* and the pumping rate was scored after 24 h. Data were normalized to MNV control worms for each experiment and expressed as the mean percentage of pumps/min ± SEM (n= 30 worms/group, 3 assays). IC_50_ values ± SEM are reported. (**d**) Worms were treated for 2 h at room temperature with 57 µM ascorbic acid, 25 µM doxycycline, or 0.5 nM PBT2 (100 worms/100 µl). Control worms were treated with 10 mM PBS, pH 7.4 (Vehicle). Worms were then plated on NGM seeded with *E. coli,* and the pumping rate was scored after 24 h. Data were normalized to MNV control worms for each experiment and expressed as the mean percentage of pumps/min ± SEM **** p<0.0001, two-way ANOVA, and Bonferroni’s *post hoc* test. Interaction MNH/MNH + ascorbic acid= p<0.0001, MNH/MNH + doxycycline= p<0.0001, MNH/MNH+ PBT2= p<0.0001.

## Discussion

We herein generated and characterized the first transgenic *C. elegans* model expressing a human amyloidogenic LC reproducing the cardiac-specific dysfunction observed in AL patients, at least in the early phases of the disease, when the amount of amyloid deposits is minimal. In particular, we expressed a λ LC, representing the most common isotype responsible for about 80% of AL cases [43].

The amyloidogenic LCs expressed in the muscle cells of MNH worms were almost wholly secreted. They reached the pharynx, causing a specific toxic effect similar to that in AL patients where LCs, secreted into the bloodstream by bone marrow plasma cells, cause cardiotoxicity and progressive severe cardiac dysfunction. No functional and structural damage was observed in the body-wall muscle tissue where LCs were produced, at least in the first week of adulthood. However, expression of amyloidogenic LC impaired MNH worms’ healthy aging process and reduced their survival.

Interestingly, the expression in MNH worms of the H7 amyloidogenic LC caused a pharyngeal defect comparable to that observed in our previous studies when the same LC was administered to wild-type *C. elegans* [21, 22]. Moreover, in the pharynx of MNH worms and in worms fed H7, there is also increased production of mitochondrial ROS and marked structural damage, resembling that found on autopsy examination of the hearts of AL patients [22].

In MNM worms, which were generated as an additional control to prove the specific effect of amyloidogenic LC expression, only a fraction of the LCs were secreted. No structural damage to the pharynx or muscle tissue was observed. The only behavioral changes were a reduction in lifespan and health span, which may be ascribed to the intracellular retention of the LC in the body-wall muscle cells of *C. elegans,* causing protein homeostasis impairment and proteotoxicity thus mimicking some pathological mechanisms possibly occurring in multiple myeloma.

We observed that the mRNA and protein levels of the non-amyloidogenic LC in MNM worms were about twice that of the amyloidogenic once expressed by MNH. This is consistent with the quality control system’s role in reducing the production of the misfolded protein [44]. Amyloidogenic LCs that escape the quality control mechanism can form dimers, unlike the non-amyloidogenic protein that forms monomers and dimers, proving the higher ability of amyloidogenic LC to form oligomers *in vivo* in *C. elegans*.

We have not found amyloid fibrils in MNH worms in the pharynx or the body-wall muscle. Based on the knowledge that protein aggregation increases with *C. elegans’* age and with the temperature to which they are exposed by regulating protein homeostasis [45, 46] and proteasome activity [45], several experiments were done on worms from the first till the nine days of adulthood, cultivating them at 20°C or 25°C. We could observe, at most, in adult worms on the nine days of adulthood maintained at 25°C an increase in the size of mCherry-positive spots in MNH::mCherry worms whose X-34 positivity was not always detectable. Given that the staining of the amyloid fibers was done by feeding the worm with the 4, due to the reduced pharyngeal functionality of MNH nematodes, the concentration of dye that arrived in conjunction with the LCs could not always be sufficient to give a specific signal. TEM analysis could not detect the presence of amyloid fibrils in the pharynx and body-wall muscle of MNH transgenic animals on the first day of adulthood. We cannot exclude that the formation of amyloid deposits occurred beyond the ten days of adulthood when worms began to die, and their pharyngeal function was so compromised as to not allow *in vivo* staining with X-34. However, the absence of amyloid deposits in MNH worms reveals that the formation of soluble oligomeric species of amyloidogenic LC can cause a toxic effect and is sufficient for disease onset.

To gain insight into the pharyngeal dysfunction of MNH worms, we registered the electropharyngeograms (i.e., the pharyngeal electrical activity), which provide information on parameters such as the rate of pharyngeal pumping and the duration of each pump [47]. We observed that the pharynx contracted less frequently in MNH worms than in control worms, possibly reflecting electrical disturbances and functional impairment occurring in AL patients’ hearts [48]. Although the circuitry of the pharynx is similar to that of the heart, with gap junctions coupling and synchronizing myocytes for potential conduction [30], it may be taken into account that many structural differences exist between proteins involved in generating potential [47].

Despite its limitations, the functionality of the nematode pharynx, similar to the mammalian heart, has been shown to decline with aging [49] and in *C. elegans* models of mitochondrioapthies [50], which often present with cardiomyopathy in humans. Notably, the MNH transgenic worms described in our work currently represent the most straightforward, least expensive, and fastest animal model to study the mechanisms underlying cardiac toxicity of LC in AL amyloidosis. Additional studies like single-cell RNA sequencing will help us understand whether organ tropism is controlled by specific genes/proteins or whether other mechanisms are involved.

As indicated by the data obtained with the prototypical drugs we used in the study, MNH worms also represent an excellent model to evaluate the preclinical efficacy of new molecules, thus offering a significant contribution to studies that could only be carried out *in vitro*, in cellular models, and using purified LCs.

## Acknowledgments

We thank Prof. Stefano Ricagno for constructive discussion and Dr. Oscar Fumagalli for his technical participation in *C. elegans* lifespan experiments. *OP50 Escherichia coli* bacteria were provided by the Caenorhabditis Genetics Center (CGC), which is funded by the National Institutes of Health (NIH) Office of Research Infrastructure Programs (P40 OD010440).

## Statements & Declarations

### Author Contributions

M.R, M.M.B, M.N., and L.D. designed research; M.R, M.M.B, A.C., S.M., C.N., A.Conz, and F.F. performed research; M.R, M.M.B, A.C., S.M., N.V., C.N., F.F., and L.D. collected data, analyzed and interpret data; M.R, M.M.B, S.M., N.V., and L.D. performed statistical analysis; M.R, M.M.B, N.V., F.F., M.S., M.N., G.P., G.M. and L.D. wrote the manuscript.

### Data availability

The article and its supplementary information contain all relevant data. All raw data have been deposited to zenodo.org and can be requested at.

### Ethics approval

Not applicable.

### Consent to participate

Not applicable.

### Consent to publish

Not applicable.

### Diversity and inclusion statement

No diversity and inclusion practices were applied to the paper’s scientific content, authorship, and attribution.

## References

1. Sabinot A, Ghetti G, Pradelli L, et al (2023) State-of-the-art review on AL amyloidosis in Western Countries: Epidemiology, health economics, risk assessment and therapeutic management of a rare disease. Blood Rev 59:101040. 10.1016/j.blre.2023.101040

2. Sanchorawala V (2024) Systemic Light Chain Amyloidosis. N Engl J Med 390:2295–2307. 10.1056/NEJMra2304088

3. Duca F, Rettl R, Kronberger C, et al (2023) Myocardial structural and functional changes in cardiac amyloidosis - Insights from a prospective observational patient registry. Eur Heart J Cardiovasc Imaging. 10.1093/ehjci/jead188

4. Lin HM, Gao X, Cooke CE, et al (2017) Disease burden of systemic light-chain amyloidosis: a systematic literature review. Curr Med Res Opin 33:1017–1031. 10.1080/03007995.2017.1297930

5. Oghina S, Delbarre MA, Poullot E, et al (2022) [Cardiac amyloidosis: State of art in 2022]. Rev Med Interne 43:537–544. 10.1016/j.revmed.2022.04.036

6. Ríos-Tamayo R, Krsnik I, Gómez-Bueno M, et al (2023) AL Amyloidosis and Multiple Myeloma: A Complex Scenario in Which Cardiac Involvement Remains the Key Prognostic Factor. Life (Basel) 13:. 10.3390/life13071518

7. Merlini G, Dispenzieri A, Sanchorawala V, et al (2018) Systemic immunoglobulin light chain amyloidosis. Nat Rev Dis Primers 4:38. 10.1038/s41572-018-0034-3

8. Palladini G, Milani P (2023) Diagnosis and Treatment of AL Amyloidosis. Drugs 83:203–216. 10.1007/s40265-022-01830-z

9. Bal S, Landau H (2021) AL amyloidosis: untangling new therapies. Hematology Am Soc Hematol Educ Program 2021:682–688. 10.1182/hematology.2021000305

10. Bou Zerdan M, Nasr L, Khalid F, et al (2023) Systemic AL amyloidosis: current approach and future direction. Oncotarget 14:384–394. 10.18632/oncotarget.28415

11. Puri S, Schulte T, Chaves-Sanjuan A, et al (2023) The Cryo-EM STRUCTURE of Renal Amyloid Fibril Suggests Structurally Homogeneous Multiorgan Aggregation in AL Amyloidosis. J Mol Biol 435:168215. 10.1016/j.jmb.2023.168215

12. Pradhan T, Sarkar R, Meighen-Berger KM, et al (2023) Mechanistic insights into the aggregation pathway of the patient-derived immunoglobulin light chain variable domain protein FOR005. Nat Commun 14:3755. 10.1038/s41467-023-39280-0

13. Fishov H, Muchtar E, Salmon-Divon M, et al (2023) AL amyloidosis clonal plasma cells are regulated by microRNAs and dependent on anti-apoptotic BCL2 family members. Cancer Med 12:8199–8210. 10.1002/cam4.5621

14. Russo R, Romeo M, Schulte T, et al (2022) Cu(II) Binding Increases the Soluble Toxicity of Amyloidogenic Light Chains. Int J Mol Sci 23:. 10.3390/ijms23020950

15. Kazman P, Absmeier RM, Engelhardt H, Buchner J (2021) Dissection of the amyloid formation pathway in AL amyloidosis. Nat Commun 12:6516. 10.1038/s41467-021-26845-0

16. Radamaker L, Karimi-Farsijani S, Andreotti G, et al (2021) Role of mutations and post-translational modifications in systemic AL amyloidosis studied by cryo-EM. Nat Commun 12:6434. 10.1038/s41467-021-26553-9

17. Swuec P, Lavatelli F, Tasaki M, et al (2019) Cryo-EM structure of cardiac amyloid fibrils from an immunoglobulin light chain AL amyloidosis patient. Nat Commun 10:1269. 10.1038/s41467-019-09133-w

18. Oberti L, Maritan M, Rognoni P, et al (2019) The concurrency of several biophysical traits links immunoglobulin light chains with toxicity in AL amyloidosis. Amyloid 26:107–108. 10.1080/13506129.2019.1583187

19. Maritan M, Ambrosetti A, Oberti L, et al (2019) Modulating the cardiotoxic behaviour of immunoglobulin light chain dimers through point mutations. Amyloid 26:105–106. 10.1080/13506129.2019.1583185

20. Zhang Y, Yu W, Chang W, et al (2023) Light Chain Amyloidosis-Induced Autophagy Is Mediated by the Foxo3a/Beclin-1 Pathway in Cardiomyocytes. Lab Invest 103:100001. 10.1016/j.labinv.2022.100001

21. Diomede L, Rognoni P, Lavatelli F, et al (2014) A Caenorhabditis elegans-based assay recognizes immunoglobulin light chains causing heart amyloidosis. Blood 123:3543–3552. 10.1182/blood-2013-10-525634

22. Diomede L, Romeo M, Rognoni P, et al (2017) Cardiac Light Chain Amyloidosis: The Role of Metal Ions in Oxidative Stress and Mitochondrial Damage. Antioxid Redox Signal 27:567–582. 10.1089/ars.2016.6848

23. Merlini G (2017) AL amyloidosis: from molecular mechanisms to targeted therapies. Hematology Am Soc Hematol Educ Program 2017:1–12. 10.1182/asheducation-2017.1.1

24. Martinez-Rivas G, Bender S, Sirac C (2022) Understanding AL amyloidosis with a little help from in vivo models. Front Immunol 13:1008449. 10.3389/fimmu.2022.1008449

25. Mishra S, Joshi S, Ward JE, et al (2019) Zebrafish model of amyloid light chain cardiotoxicity: regeneration versus degeneration. Am J Physiol Heart Circ Physiol 316:H1158–H1166. 10.1152/ajpheart.00788.2018

26. Ayala MV, Bender S, Anegon I, et al (2021) A rat model expressing a human amyloidogenic kappa light chain. Amyloid 28:209–210. 10.1080/13506129.2021.1877651

27. Nuvolone M, Sorce S, Pelczar P, et al (2017) Regulated expression of amyloidogenic immunoglobulin light chains in mice. Amyloid 24:52–53. 10.1080/13506129.2017.1289914

28. Martinez-Rivas G, Ayala M, Bender S, et al (2024) A mouse model of cardiac AL amyloidosis unveils mechanisms of tissue accumulation and toxicity of amyloid fibrils. bioRxiv 2024.07.18.604040. 10.1101/2024.07.18.604040

29. Benian GM, Epstein HF (2011) Caenorhabditis elegans muscle: a genetic and molecular model for protein interactions in the heart. Circ Res 109:1082–1095. 10.1161/CIRCRESAHA.110.237685

30. Mango SE (2007) The C. elegans pharynx: a model for organogenesis. WormBook 1–26. 10.1895/wormbook.1.129.1

31. Avery L, Shtonda BB (2003) Food transport in the C. elegans pharynx. J Exp Biol 206:2441–2457. 10.1242/jeb.00433

32. Jones D, Candido EP (1999) Feeding is inhibited by sublethal concentrations of toxicants and by heat stress in the nematode Caenorhabditis elegans: relationship to the cellular stress response. J Exp Zool 284:147–157. 10.1002/(sici)1097-010x(19990701)284:2<147::aid-jez4>3.3.co;2-q

33. Haun C, Alexander J, Stainier DY, Okkema PG (1998) Rescue of Caenorhabditis elegans pharyngeal development by a vertebrate heart specification gene. Proc Natl Acad Sci U S A 95:5072–5075. 10.1073/pnas.95.9.5072

34. Nas JSB (2021) Caenorhabditis elegans as a Model in Studying Physiological Changes Following Heart Failure. Asian Journal of Biological and Life Sciences, 10:522–6. 10.5530/ajbls.2021.10.69

35. Frøkjær-Jensen C, Wayne Davis M, Hopkins CE, et al (2008) Single-copy insertion of transgenes in Caenorhabditis elegans. Nature Genetics 40:1375–1383. 10.1038/ng.248

36. Pir GJ, Choudhary B, Mandelkow E, Mandelkow E-M (2016) Tau mutant A152T, a risk factor for FTD/PSP, induces neuronal dysfunction and reduced lifespan independently of aggregation in a C. elegans Tauopathy model. Mol Neurodegener 11:33. 10.1186/s13024-016-0096-1

37. Tahseen Q (2009) Coelomocytes: Biology and Possible Immune Functions in Invertebrates with Special Remarks on Nematodes. In: International Journal of Zoology. https://www.hindawi.com/journals/ijz/2009/218197/. Accessed 1 Aug 2023

38. Egami Y, Taguchi T, Maekawa M, et al (2014) Small GTPases and phosphoinositides in the regulatory mechanisms of macropinosome formation and maturation. Front Physiol 5:374. 10.3389/fphys.2014.00374

39. Styren SD, Hamilton RL, Styren GC, Klunk WE (2000) X-34, a fluorescent derivative of Congo red: a novel histochemical stain for Alzheimer’s disease pathology. J Histochem Cytochem 48:1223–1232. 10.1177/002215540004800906

40. Link CD, Johnson CJ, Fonte V, et al (2001) Visualization of fibrillar amyloid deposits in living, transgenic Caenorhabditis elegans animals using the sensitive amyloid dye, X-34. Neurobiol Aging 22:217–226. 10.1016/s0197-4580(00)00237-2

41. Ward JE, Ren R, Toraldo G, et al (2011) Doxycycline reduces fibril formation in a transgenic mouse model of AL amyloidosis. Blood 118:6610–6617. 10.1182/blood-2011-04-351643

42. Summers KL, Roseman G, Schilling KM, et al (2022) Alzheimer’s Drug PBT2 Interacts with the Amyloid β 1-42 Peptide Differently than Other 8-Hydroxyquinoline Chelating Drugs. Inorg Chem 61:14626–14640. 10.1021/acs.inorgchem.2c01694

43. Desport E, Bridoux F, Sirac C, et al (2012) Al amyloidosis. Orphanet J Rare Dis 7:54. 10.1186/1750-1172-7-54

44. Diomede L, Soria C, Romeo M, et al (2012) C. elegans expressing human β2-microglobulin: a novel model for studying the relationship between the molecular assembly and the toxic phenotype. PLoS One 7:e52314. 10.1371/journal.pone.0052314

45. Lee HJ, Alirzayeva H, Koyuncu S, et al (2023) Cold temperature extends longevity and prevents disease-related protein aggregation through PA28γ-induced proteasomes. Nat Aging 3:546–566. 10.1038/s43587-023-00383-4

46. Cuanalo-Contreras K, Schulz J, Mukherjee A, et al (2022) Extensive accumulation of misfolded protein aggregates during natural aging and senescence. Front Aging Neurosci 14:1090109. 10.3389/fnagi.2022.1090109

47. Raizen DM, Avery L (1994) Electrical activity and behavior in the pharynx of Caenorhabditis elegans. Neuron 12:483–495. 10.1016/0896-6273(94)90207-0

48. Falk RH, Alexander KM, Liao R, Dorbala S (2016) AL (Light-Chain) Cardiac Amyloidosis: A Review of Diagnosis and Therapy. J Am Coll Cardiol 68:1323–1341. 10.1016/j.jacc.2016.06.053

49. Eckers A, Jakob S, Heiss C, et al (2016) The aryl hydrocarbon receptor promotes aging phenotypes across species. Sci Rep 6:19618. 10.1038/srep19618

50. Maglioni S, Schiavi A, Melcher M, et al (2022) Neuroligin-mediated neurodevelopmental defects are induced by mitochondrial dysfunction and prevented by lutein in C. elegans. Nat Commun 13:2620. 10.1038/s41467-022-29972-4

51. Perfetti V, Sassano M, Ubbiali P, et al (1996) Inverse polymerase chain reaction for cloning complete human immunoglobulin variable regions and leaders conserving the original sequence. Anal Biochem 239:107–109. 10.1006/abio.1996.0297

52. Garofalo M, Piccoli L, Romeo M, et al (2021) Machine learning analyses of antibody somatic mutations predict immunoglobulin light chain toxicity. Nat Commun 12:3532. 10.1038/s41467-021-23880-9

53. Zanier ER, Barzago MM, Vegliante G, et al (2021) C. elegans detects toxicity of traumatic brain injury generated tau. Neurobiol Dis 153:105330. 10.1016/j.nbd.2021.105330

54. Lockery SR, Hulme SE, Roberts WM, et al (2012) A microfluidic device for whole-animal drug screening using electrophysiological measures in the nematode C. elegans. Lab Chip 12:2211–2220. 10.1039/c2lc00001f

55. Han SK, Lee D, Lee H, et al (2016) OASIS 2: online application for survival analysis 2 with features for the analysis of maximal lifespan and healthspan in aging research. Oncotarget 7:56147–56152. 10.18632/oncotarget.11269

