## Supplemental Materials, Tables and Figures for "Modeling immunoglobulin light chain amyloidosis in *Caenorhabditis elegans*"

**Methods**

**mRNA extraction, PCR, and quantitative PCR (Q-PCR)**

The nematodes were synchronized by egg-laying and cultured on NGM plates seeded with *E. coli* OP50 for food at 20°C. On the first day of adulthood, nematodes were collected with M9 buffer, settled by gravity, and washed with M9 to eliminate bacteria. According to the manufacturer's instructions, RNA was extracted from the pellet of worms using the Maxwell® RSC simplyRNA Tissue kit (Promega Italia Srl, Milan, Italy). Briefly, the pellet of worms was homogenized by Turrax (T10, IKA-Werke GmbH & Co., Germany) for 2 min using 200 μl of a chilled working solution prepared by adding 20 μl of 1-thioglycerol per milliliter of homogenization solution. Lysis buffer (200 μl) was added to 200 μl of homogenate, vortexed for 15 seconds, and transferred into the cartridge well. RNA was extracted and eluted in 50 μl of nuclease-free water, and the concentration was quantified using NanoDrop (Thermo Fisher Scientific Inc., Monza, Italy). mRNA was reverse transcribed using the FIREScript RT cDNA Synthesis kit (Solis BioDyne, Tartu, Estonia). To this end, 1 µg of RNA was retrotranscripted using oligo (dT) primer in a 20 µl final mix volume. cDNA (50 ng) was used for Q-PCR amplification using the 2X Power SYBR^TM^ Green PCR Master mix (Applied Biosystem, Thermo Fisher Scientific, MA, USA) and the Applied Biosystem 7300 Real-Time PCR System. Two specific primers were designed on the constant region of H7 and M7 sequences and were used for the PCR amplification (**Supplementary Table 2).** The relative level of H7 and M7 genes expressed by MNH and MNM worms were determined using the 2^-ΔΔCT^ method using the cell division cycle-related (*cdc-42*) and the conserved iron-binding-related (*y45f10d.4*) as housekeeping genes (**Supplementary Table 2)**.

**Western blot analysis**

The nematodes were synchronized by egg-laying and cultured on NGM plates seeded with *E. coli* OP50 for food at 20°C. On the first day of adulthood, nematodes were collected with M9 buffer, settled by gravity, and washed with M9 to eliminate bacteria. To evaluate the total LC protein levels, pellets from 50 worms were suspended in 50 μl 1X SDS loading sample buffer (10% glycerol, 2% SDS, 60 mM Tris, pH 6.8) added with 5% β-mercaptoethanol. Samples were boiled for 10 minutes at 95°C, and 25 μl were loaded into 15% SDS-Page Gel wells. Recombinant H7 protein (50 ng), kindly provided by Prof. Stefano Ricagno (University of Milan, Milan, Italy), was loaded as a control. To perform Western blot analysis under non-reducing conditions, the worm pellets were suspended in 200 μl cold 10 mM PBS supplemented with complete protease inhibitors (Roche, Basel, Switzerland) and homogenized in ice by Turrax homogenization for 2 min. Samples were spun at 5900 x *g* for 5 minutes at 4°C, the supernatant was collected, and the protein concentration was determined using Pierce BCA Protein Assay Kit (Thermo Fisher Inc.). Samples (25 μg total protein) were suspended in 1X SDS loading sample buffer without β-mercaptoethanol, boiled for 5 minutes at 95°C, and loaded into the wells of a 15% SDS-Page Gel. Recombinant H7 (50 ng) was loaded alone or with 25 µg of protein lysate of MNV worms as positive controls. To quantify secreted and non-secreted LCs, the worm pellets were suspended in 200 μl cold 10 mM PBS supplemented with complete protease inhibitors (Roche, Basel, Switzerland) and homogenized in ice by Turrax homogenization for 2 min. Samples were spun at 3000 x *g* for 15 minutes at 4°C, and the supernatant was collected as the fraction containing secreted LCs. The pellet, collected as the fraction containing the non-secreted LCs [1], was washed twice with 10 mM PBS, re-suspended in a hot 10% SDS solution, and boiled for 10 minutes at 95°C. The protein concentration of secreted and non-secreted fractions was determined using the Pierce BCA Protein Assay Kit. Twenty-five μg of total proteins were suspended in 1X SDS loading sample buffer containing 5% β-mercaptoethanol, boiled for 5 minutes at 95°C, and loaded into the wells of a 15% SDS-Page Gel.

To assess the LC solubility, proteins were extracted from transgenic worms on the first day of adulthood using RAB (100 mM 2-(N-morpholino) ethanesulfonic acid, 1 mM EGTA, 0.5 mM MgSO_4_, 20 mM NaF) and RIPA (150 mM NaCl, 1% (v/v) Nonidet P-40, 0.5% (w/v) deoxycholate, 0.1% sodium dodecyl sulfate, 50 mM, Tris, pH 8.0) buffers containing EDTA-free protease inhibitors as already described [2].

Proteins in all the SDS-Page gels were separated at 100 V in SDS running buffer and blotted for 2 h at 100V onto a PVDF membrane in 20 mM Tris solution containing 150 mM glycine and 10% methanol. Membranes were blocked for 1 h at room temperature in 10 mM Tris-HCl solution, pH 7.5, containing 100 mM NaCl, 0.1% (v/v) Tween 20, 5% (w/v) low-fat dry milk powder, and 2% (w/v) bovine serum albumin, and incubated overnight at 4°C with a mouse monoclonal antibody anti-human λ light chains (Bound and Free) (1:1000 dilution, Sigma-Aldrich, Milan, Italy), or a mouse monoclonal anti-actin antibody clone C4 (1:2000 dilution, Sigma-Aldrich). Anti-mouse IgG peroxidase conjugate (1:10000, Sigma-Aldrich) was used as the secondary antibody. Chemioluminescence was detected by Clarity Max Western ECL Substrate (Biorad, Hercules, California, USA), and the membranes were scanned with a ChemiDoc Imaging System (Biorad). Chemioluminescence was detected by Clarity Max Western ECL Substrate (Biorad, Hercules, California, USA), and the membrane images were acquired with the ChemiDoc Imaging System (Biorad). The mean volumes of immunoreactive bands were determined using Image Lab™ software (Bio-Rad).

**Lifespan and healthspan**

The nematodes were synchronized by egg-laying and transferred to NGM plates daily during the fertile period to avoid overlapping generations. Dead, alive, and censored animals were scored during the transferring process. The animals were counted as dead when they had neither moved nor reacted to a manual stimulus with a platinum wire nor had any pharyngeal pumping activity. Animals with exploded vulvas or those desiccated on the wall were censored [3]. The number of active movements was also assessed in nematodes employed for the lifespan assay to determine their healthy aging. Animal crawling spontaneously or after a manual stimulus was considered moving, while dead animals and animals without crawling behavior were considered not moving.

**Mitochondrial ROS production**

Mitochondrial ROS production was detected in live worms using MitoSOX™ Red staining (ThermoFisher Scientific, Milan, Italy). The nematodes were synchronized by egg-laying and cultivated on NGM plates until the L4 larval stage. Sixty worms were transferred onto freshly prepared 6 cm NGM plates containing 10 µM MitoSOX™ Red and seeded with UV-killed *E. Coli* OP50 bacteria. After incubating for 16 h in the dark at 20°C, when worms were on the first day of adulthood, they were transferred for 1 h to new NGM plates spread with live OP50 bacteria to remove residual dye from the gut. As positive control, MNV worms at the L4 larval stage were incubated for 2 hours with 0.5 mM H_2_O_2_, plated onto 6 cm NGM plates containing 10 µM MitoSOX™ Red, and processed as described before.

For imaging, nematodes were mounted onto 2% agarose pad slides, anesthetized by adding 10 mM levamisole (Sigma-Aldrich), and fixed by ProLong™ Glass Antifade Mountant (ThermoFisher Scientific). Images were acquired immediately with an inverted fluorescent microscope (IX-71 Olympus, Tokyo, Japan) equipped with a CCD camera. Pictures of the pharynx were obtained at 40X magnification with a TRITC filter set (Olympus), and the integrated intensity was calculated using Fiji’s imaging software [4].

**Fluorescent and confocal microscopy**

Age-synchronized MNH::mCherry worms were picked onto 2% agar pads, anesthetized with 10 mM levamisole, and fixed by ProLong™ Glass Antifade Mountant on the first day of adulthood. Images were acquired with an inverted fluorescent microscope (IX-71 Olympus) equipped with a CCD camera. Pictures of the pharynges were taken at 40× magnification with a TRITC filter set (Olympus). Images were also acquired by confocal microscopy using Nikon A1 Confocal microscopes (Nikon).

**TABLES**

**Supplementary Table 1.** **List of the strains used in this study**

| Strain name abbreviation | Strain | Genotype |
| --- | --- | --- |
| MNH | COP2040 - knuSi825 | *[pNU2100 (myo-3::H7::tbb-2u in ttTi5605,unc-119(+) ) ] II ; unc-119(ed3) III* |
| MNM | COP2177 - knuSi837 | *[pNU2101(myo-3::M7::tbb-2u in ttTi5605, unc-119(+) ) ] II; unc-119(ed3) III* |
| MNV | COP2043 - knuSi827 | *[pNU936(empty ttTi5605,*  *unc-119(+))] II ; unc-119(ed3) III* |
| MNH::mCherry | COP2459 - knuls85 | *[myo-3p::H7::mCherry::tbb-2 3’-UTR * knuSi825) ] II* |

**Supplementary Table 2. List of the primers used in this study**

| **Name** | **Sequence (5’-3’)** |
| --- | --- |
| H7/M7-Constant region- Forward | 5’-CTCTTCCCACCATCCTCC-3’ |
| H7/M7-Constant region -Reverse | 5’-CGGCGTACTTGTTGTTGGAT-3’ |
| cdc-42 -Forward | 5’-CTGTTGTGGTGGGTCGAGAG-3’ |
| cdc-42 -Reverse | 5’-GTTGACGCAGAAGGGACTGA-3’ |
| Y45F10D.4 -Forward | 5’-ATCTTCCCTGGCAACCGAAT-3’ |
| Y45F10D.4 -Reverse | 5’-TGGGCGAGCATTGAACAGT-3’ |

**Supplementary Table 3. Summary of lifespan and health span analysis**

|  | **Strain** | **MEAN**  **(Days ± SEM)** | **p-value**  ***vs***  **MNV** | **p-value**  ***vs***  **MNH** | **No. of subjects** | **Censor** |
| --- | --- | --- | --- | --- | --- | --- |
| **Lifespan** | **MNV** | 18.9 **±** 0.5 |  | 0.0001 | 180 | 21 |
|  | **MNH** | 16.7**±** 0.3 | 0.0001 |  | 180 | 23 |
|  | **MNM** | 15.6**±** 0.3 | <0.0001 | 0.0044 | 180 | 38 |
| **Healthspan** | **MNV** | 16.1**±** 0.4 |  | 0.0081 | 180 | 21 |
|  | **MNH** | 15.1**±** 0.3 | 0.0081 |  | 180 | 23 |
|  | **MNM** | 14.0**±** 0.2 | <0.0001 | 0.0016 | 180 | 38 |

The p-values were calculated using the log-rank and Bonferroni’s *post hoc* test between the pooled populations of animals.

**FIGURES**

**H7 Sequence**

ATG**ACCTGCTCCCCTCTCCTCCTCACCCTCCTTATTCACTGCACCGGATCCTGGACC**CAATCCGTCCTCACCCAACCACCATCCGTCTCCGCCGCCCCAGGACAAAAGGTCACCATCTCCTGCTCCAACGTCGGAAAGAACTTCGTCTCCTGGTACCAACAATTCCCAGGAACCGCCCCAAAGGTCGTCATCTACGACACCGACAAGCGTCCATCCGACATCCCAGACCGTTTCTCCGGATCCAAGTCCGGAACCTCCGCCACCCTCGACATCACCGGACTCCAAACCGGAGACGAGGCCGACTACTACTGCGGAACCTGGGACTCCGGACTCAACGGAGGAGTCTTCGGAGGAGGAACCAAG**gtaagtttaaacatatatatactaactaaccctgattatttaaattttcag**GTCACCGTCCTCGGACAACCAAAGGCCGCCCCATCCGTCACCCTCTTCCCACCATCCTCCGAGGAGCTCCAAGCCAACAAGGCCACCCTCGTCTGCCTCATCTCCGACTTCTACCCAGGAGCCGTCACCGTCGCCTGGAAGGCCGACTCCTCCCCAGTCAAGGCCGGAGTCGAGACCACCACCCCATCCAAGCAATCCAACAACAAGTACGCCGCCTCCTCCTACCTCTCCCTCACCCCAGAGCAATGGAAGTCCCACAAGTCCTACTCCTGCCAAGTCACCCACGAGGGATCCACCGTCGAGAAGACCGTCGCCCCAACCGAGTGCTCCTAA

**M7 sequence**

ATG**GCTTGGACTCCTTTATGGCTCACTCTCCTTACGCTGTGCATCGGATCCGTCGTC**TCCTCCGAGCTCACCCAAGACCCAGCCGTCTCCGTCGCCCTCGGACAAACCGTCAAGATCACCTGCCAAGGAGACTCCCTCCGTATGTACTACGCCTCCTGGTACCAACAAAAGCCAGCCCAAGCCCCAGTCCTCGTCATCTACGCCGAGAAGAACCGTCCATCCGGAATCCCAGACCGTTTCTCCGCCTCCTCCTCCGGATCCACCGCCTCCCTCACCATCACCGGAGCCCAAGCCGAGGACGAGGCCGACTACTACTGCAACTCCCGTGACAACTCCGGAGACCACCTCGTCTTCGGAGGAGGAACCAAGCTCACCGTCCTCGGACAACCAAAG**gtaagtttaaacatatatatactaactaaccctgattatttaaattttcag**GCCGCCCCATCCGTCACCCTCTTCCCACCATCCTCCGAGGAGCTCCAAGCCAACAAGGCCACCCTCGTCTGCCTCATCTCCGACTTCTACCCAGGAGCCGTCACCGTCGCCTGGAAGGCCGACTCCTCCCCAGTCAAGGCCGGAGTCGAGACCACCACCCCATCCAAGCAATCCAACAACAAGTACGCCGCCTCCTCCTACCTCTCCCTCACCCCAGAGCAATGGAAGTCCCACCGTTCCTACTCCTGCCAAGTCACCCACGAGGGATCCACCGTCGAGAAGACCGTCGCCCCAACCGAGTGCTCCTAA

**Supplementary Fig. 1 The DNA sequence of the amyloidogenic cardiotoxic H7 LC and the non-amyloidogenic myeloma-derived M7 LC optimized to be expressed in *C. elegans*.**

The human DNA secretion sequence is reported in red, and the intron sequence is written in blue.

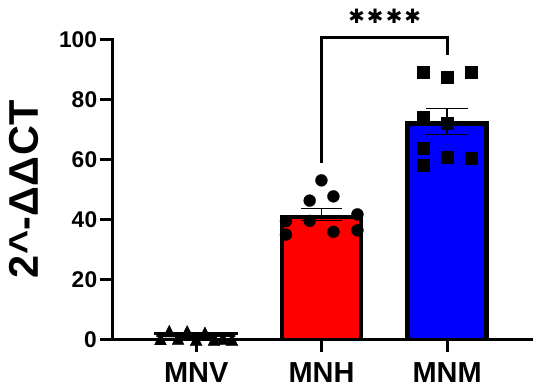

**Supplementary Fig. 2 Messenger RNA levels in transgenic worms**

Quantitative real-time PCR analysis of mRNA levels in synchronized nematodes on the first day of adulthood. Specific primers recognizing the constant LC region were used. LC gene expression levels were compared to *cdc-42* and *y45f10d.4* housekeeping mRNA. Data are the mean ± SEM (n=9, from 3 different biological samples). ****p<0.0001 according to one-way ANOVA and Bonferroni's *post hoc* test.

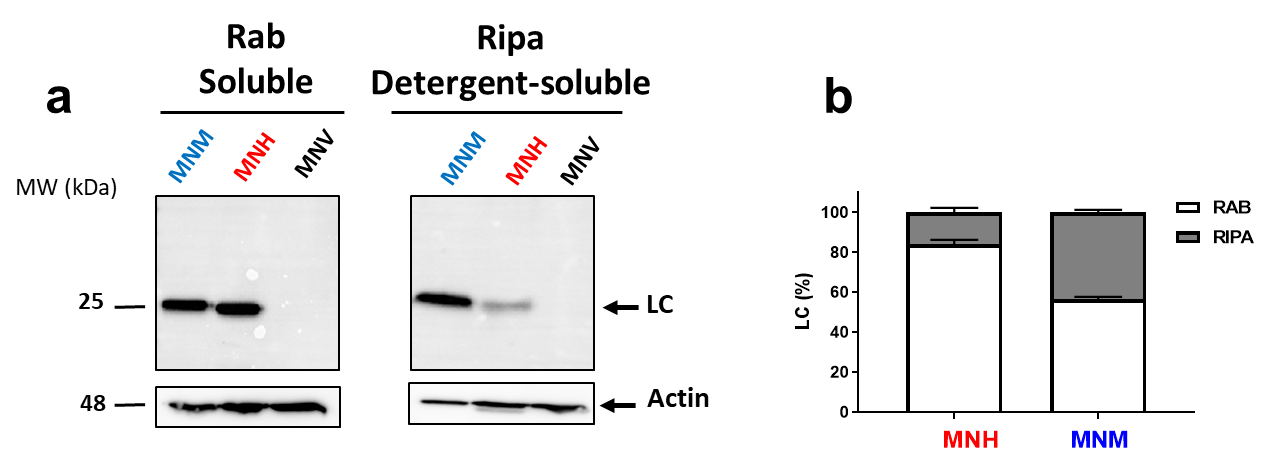

**Supplemental Fig. 3 Levels of the RAB soluble and RIPA detergent-soluble LCs**

(**a**) RAB soluble and RIPA detergent-soluble fractions from the equivalent amounts of proteins from worms on the first day of adulthood were analyzed by immunoblotting using the anti-LC antibody (1:1000 dilution) or anti-actin antibody (1:2000 dilution). (**b**) LC/actin immunoreactivity in each fraction was determined and expressed as a percentage of the total LC. Data are mean ± SD (n = 3 from 2 assays). The LC percentage in RAB soluble fraction was higher in MNH worms (84%) compared to MNM (56%). The percentage of LC in the RIPA fraction was 16% and 43% in MNH and MNM, respectively.

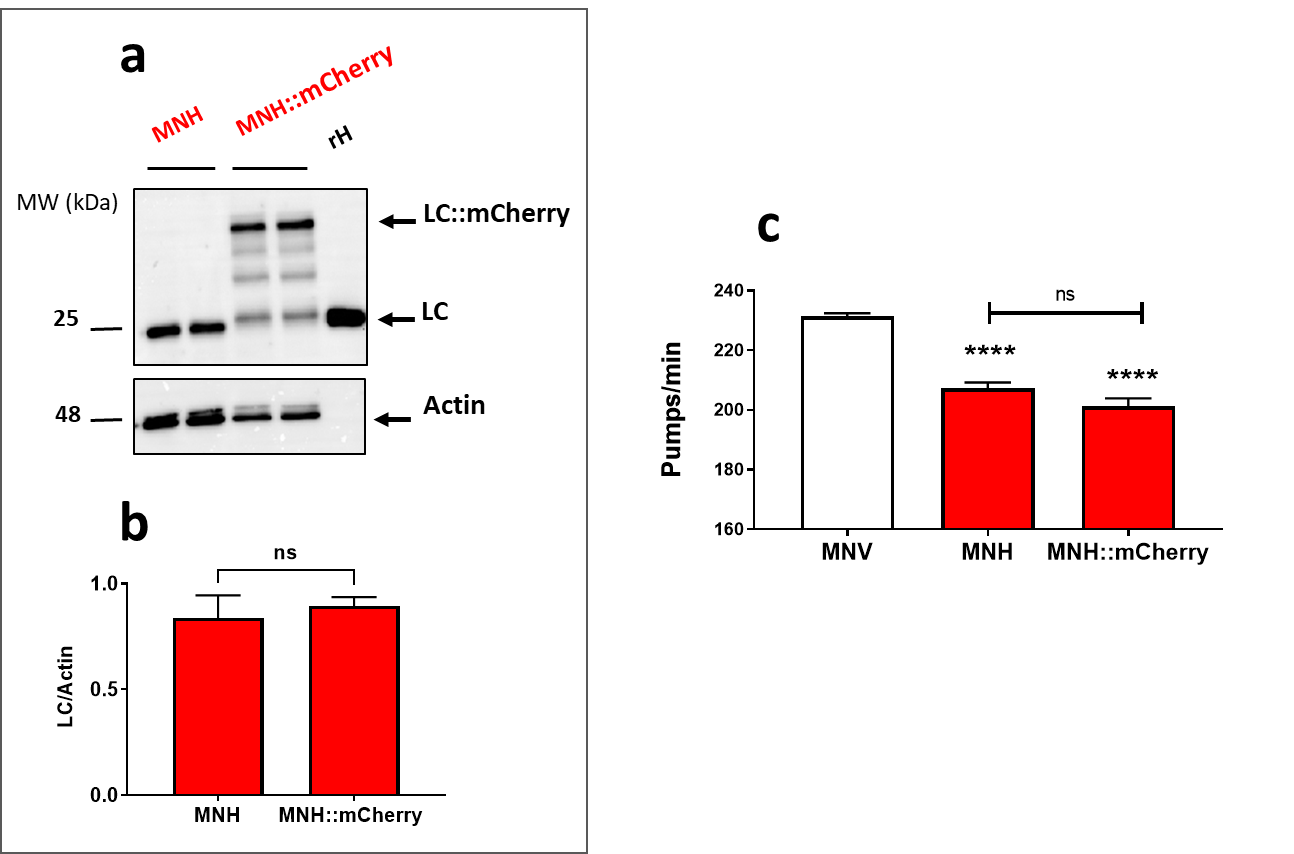

**Supplementary Fig. 4 Characterization of the transgenic** **MNH::mcherry *C. elegans* strain.**

(**a**) Representative Western blot of LC in lysates of MNH and MNH::mcherry worms on the first day of adulthood. An equal amount of proteins (25 µg) were loaded in each gel lane and immunoblotted with anti-human λ total LC or anti-actin antibody. Recombinant H7 LC (rH, 50 ng) was loaded as control. (**b**) Quantification of total LC expressed as the mean volume of the anti-human λ total LC band immunoreactivity of the Western blot in the (a)/actin band. Data are mean ± SEM (n = 3, from 2 independent experiments). No statistical difference (Student’s t-test, p= 0.1997) was found between MNH and MNH::mCherry strains. (**c**) The pharyngeal activity of worms on the first day of adulthood is expressed as pumps/min. Data are the mean ± SEM (n = 30 worms/assay, four assays). **** p<0.0001 *vs.* MNV, one-way ANOVA, and Bonferroni’s *post hoc* test.

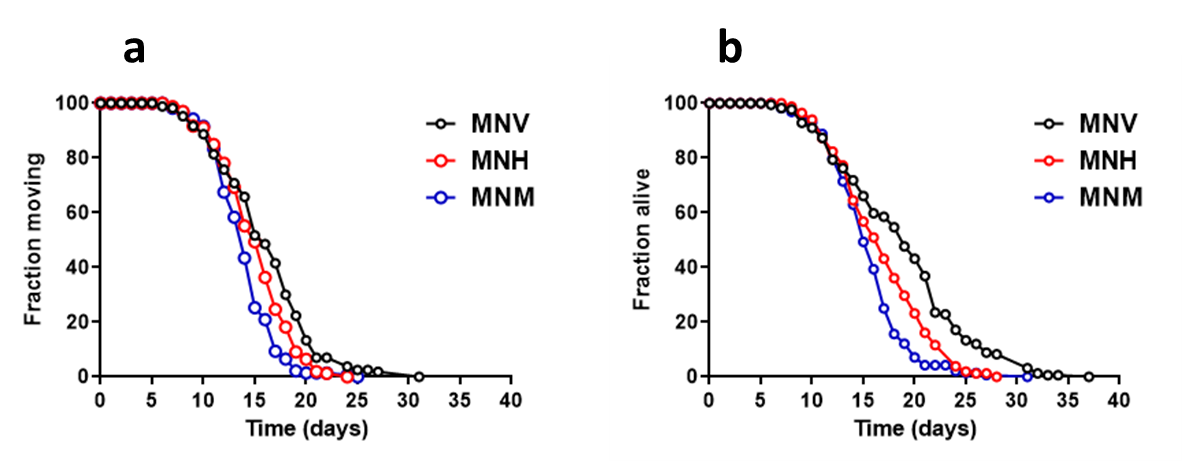

**Supplementary Fig. 5 Health span and lifespan of transgenic worms**

(**a**) Healthspan and (**b**) lifespan curves of MNV, MNH, and MNM nematodes. Dead, alive, and censored animals were scored. Data are the mean ± SEM (n=180 worms from 3 independent experiments). See Supplemental Table 3 for mean lifespan, health span, and statistical analyses.

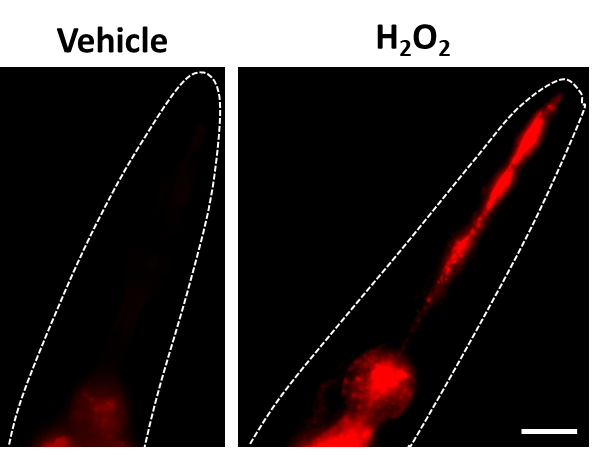

**Supplementary Fig. 6 Effect of hydrogen peroxide on mitochondrial superoxide production** Representative images of mitochondrial superoxide production in the pharynx of MNV worms administered with 10 mM PBS (Vehicle) or 0.5 mM H_2_O_2_ (H_2_O_2_), detected with MitoSOX™ Red. Scale bar = 50 µm.

**References**

1.	Madhivanan K, Greiner ER, Alves-Ferreira M, et al (2018) Cellular clearance of circulating transthyretin decreases cell-nonautonomous proteotoxicity in *Caenorhabditis elegans*. Proceedings of the National Academy of Sciences 115:E7710–E7719. https://doi.org/10.1073/pnas.1801117115

2.	Morelli F, Romeo M, Barzago MM, et al (2018) V363I and V363A mutated tau affect aggregation and neuronal dysfunction differently in C. elegans. Neurobiology of Disease 117:226–234. https://doi.org/10.1016/j.nbd.2018.06.018

3.	Leiser SF, Jafari G, Primitivo M, et al (2016) Age-associated vulval integrity is an important marker of nematode healthspan. Age (Dordr) 38:419–431. https://doi.org/10.1007/s11357-016-9936-8

4.	Brinkmann V, Romeo M, Larigot L, et al (2022) Aryl Hydrocarbon Receptor-Dependent and -Independent Pathways Mediate Curcumin Anti-Aging Effects. Antioxidants (Basel) 11:. https://doi.org/10.3390/antiox11040613
